# Mechanically Sensitive HSF1 is a Key Regulator of Left-Right Symmetry Breaking in Zebrafish Embryos

**DOI:** 10.1101/2021.02.25.432863

**Authors:** Jing Du, Shu-Kai Li, Liu-Yuan Guan, Zheng Guo, Jiang-Fan Yin, Li Gao, Toru Kawanishi, Atsuko Shimada, Qiu-Ping Zhang, Li-Sha Zheng, Yi-Yao Liu, Xi-Qiao Feng, Dong-Yan Chen, Hiroyuki Takeda, Yu-Bo Fan

## Abstract

The left-right symmetry breaking of vertebrate embryos requires fluid flow (called nodal flow in zebrafish). However, the molecular mechanisms that mediate the asymmetric gene expression regulation under nodal flow remain elusive. In this paper, we report that heat shock factor 1 (HSF1) is asymmetrically activated in the Kuppfer’s vesicle at the early stage of zebrafish embryos in the presence of nodal flow. Deficiency in HSF1 expression caused a significant *situs inversus* and disrupted gene expression asymmetry of nodal signaling proteins in zebrafish embryos. Further studies demonstrated that HSF1 could be immediately activated by fluid shear stress. The mechanical sensation ability of HSF1 is conserved in a variety of mechanical stimuli in different cell types. Moreover, cilia and the Ca^2+^-Akt signaling axis are essential for the activation of HSF1 under mechanical stress *in vitro* and *in vivo*. Considering the conserved expression of HSF1 in organisms, these findings unveil a fundamental mechanism of gene expression regulation triggered by mechanical clues during embryonic development and other physiological and pathological transformations.

## Introduction

Biological flow is necessary for vertebrate organogenesis, neuron migration, cardiovascular development, and left-right (LR) symmetry breaking during embryogenesis^1^. In vertebrates, the surface of the body is symmetrical, but the internal organs and tissues are asymmetrical. Such a development process is regulated by left-right organizers (LROs). The zebrafish LRO, Kupfer’s vesicle (KV), is a transient, fluid-filled vesicle organ. The rotation of primary cilia in KV produces counterclockwise directional nodal flow, which is necessary and sufficient for breaking left-right symmetry^2,3^. In KV, how the organism perceives node flow and the mechanism of symmetry breaking regulation are yet unclear. Two possibilities have been proposed: the chemical and mechanical senses of cilia. In the ciliary chemosensory hypothesis, nodal flow may carry a morphogen, a signaling molecule, which is released into the flow and is accepted by the cilia’s chemosensing function^4^. On the left and right sides of KV, the difference in the morphogenesis concentration detected by the receptor should be the determinant to destroy the left-right symmetry. To date, however, no such morphogen molecule has been detected, although previous studies have demonstrated the presence of chemical sensations in motile cilia^5^. On the other hand, the mechanosensory cilia that can sense the flow of fluid are involved in the occurrence of polycystic kidney disease^6^. Moreover, the nodal flow velocities on the left and right sides of zebrafish KV are not the same^7,8^. This finding indicates the possibility for the mechanical regulation function of cilia in the left-right symmetry breaking. However, it remains unclear how the difference in the mechanical microenvironment in KV is sensed by cilia and transduced into gene expression regulation. In the left-right organizers, mechanical and chemical senses are not necessarily mutually exclusive and but most likely coupled in a synergistic manner in zebrafish embryos. Although several signal molecules, such as Ca^2+^ and β-catenin, play important roles in the development of LR asymmetry^9,10^, the signal transduction mechanisms mediating nodal flow to gene expression regulation are largely unknown.

Regarding to gene transcriptional regulation, the heat shock protein (HSP) family of molecular chaperones shows a rapid and large transcriptional increase in response to diverse stresses, such as thermal and oxidative stress, indicating that these proteins are part of a fundamental defense against stresses^11-13^. The master regulator of the heat shock response is heat shock factor 1 (HSF1), the key transcriptional activator of HSPs^13^. HSF1 is a highly conserved transcription factor in organisms. In addition to HSPs, HSF1 also regulates (induces or represses) the expression of other genes involved in a wide range of cellular behaviors, including apoptosis, protein trafficking, and energy generation^14-17^. Notably, the expression of FOS and JUN, two famous and well-characterized immediate-early genes, is induced in heat-shocked cells by HSF1^18,19^. Thus, the rapid activation of HSF1 has served as a model for immediate transcriptional responses and has fostered many ground-breaking insights into the regulatory mechanisms of gene expression.

In this paper, the function of HSF1 in the LR symmetry breaking process under nodal flow was addressed in zebrafish embryos. By combining in vivo and in vitro experiments, we found that HSF1 was mechanically sensitive and immediately activated by fluid shear stress and other types of mechanical stimuli though cilia, which was pivotal for the LR axis establishment during zebrafish embryonic development.

## Results

### 1. HSF1 is asymmetrically activated in Kupffer’s vesicle of zebrafish embryos

Given that HSF1 is widely expressed throughout the early development of zebrafish embryos and conversely responds to acute stresses such as heat shock stress^20,21^, we examined the expression and activity of HSF1 in KV during LR axis determination of zebrafish embryos. We found that hsf1 and hsf2 mRNA expressions were observed at the 5-8 somite stage (Figure 1a). Interestingly, as the hallmark of HSF1 activation^22^, HSF1 phosphorylation at serine 326 was significantly elevated on the left side of KV in zebrafish embryos at the 8-somite stage (Figure 1b and c). HSF1 target gene hsp70 mRNA *in situ* hybridization also demonstrated left-biased expression of hsp70 in KV, further indicating asymmetric activation of HSF1 (Figure 1d and e). To investigate the requirement of nodal flow in the left-biased activation of HSF1 in KV, the c21orf59 gene was knocked down using a morpholino (MO), which impaired cilia mobility and disrupted nodal flow^23^. The results showed that compared with WT and control morpholino embryos, c21orf59 morphants showed comparable HSF1 activity between the left and right sides of KV (Figure 1f and g), indicating that the asymmetric activation of HSF1 is dependent on nodal flow. These observations suggest that HSF1 is asymmetrically activated in the KV of zebrafish embryos under the action of nodal flow.

**Figure 1.**
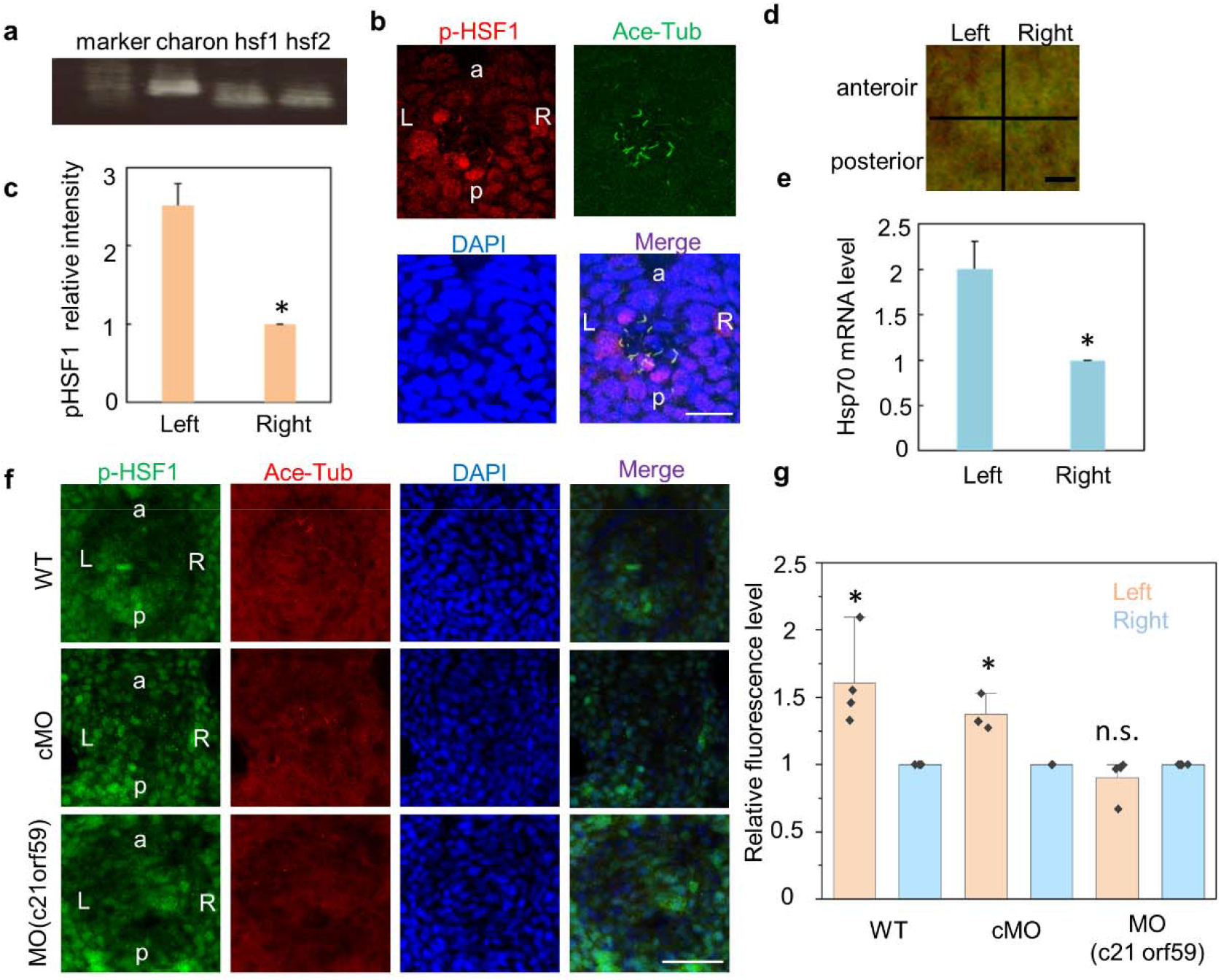
HSF1 is asymmetrically activated in KV of zebrafish embryos. (a) Expression of charon, hsf-1 and hsf-2 was examined by RT-PCR using cDNA from whole embryos of 5-8 somite stage zebrafish. (b) Immunostaining of phosphorylated HSF1 at Serine 326 and acetylated α-tubulin in KV of zebrafish embryos at 8-somite stage. Scale bar: 25μm. (c) Statistical analysis of the expression level of pS326 HSF1 in the left and right part of KV in (a) (n = 4). (d) Whole mount in situ hybridization of hsp70 mRNA in KV of zebrafish embryos at 10-somite stage. Scale bar: 10 μm. (e) Statistical analysis of the expression level of hsp70 mRNA in the left and right part of KV in (d) (n = 7). (f) Immunostaining of wild type zebrafish embryos (WT) or embryos treated with c21orf59 MO or control morpholino (cMO). Blue-DAPI, Green-p-HSF1, Red-Ace-tubulin. Scale bar: 50μm. (g) Statistical analysis of the expression level of pS326 HSF1 in the left and right part of KV in (f) (n=4)

### 2. HSF1 is required for the establishment of LR asymmetry in zebrafish embryos

Considering the important roles of HSF1 in gene expression regulation, we sought to evaluate the function of HSF1 in the LR axis establishment of zebrafish embryos and gene expression patterning asymmetry in KV. Morpholino knockdown was performed to inhibit the expression of HSF1. As shown in Figure 2a-c, injection of HSF1 morpholino into zebrafish embryos at the one- to eight-cell stage resulted in a severely randomized localization of viscera (indicated by cardiac primordia, 52% of the HSF1 MO and 8% of control MO). Moreover, supplementation with hsf1 mRNA largely rescued the aberrance in HSF1 morphants (Figure 2a-c). The effect of HSF1 MO was confirmed by CRISPR/Cas9-mediated knockout of hsf1, which caused 2-fold embryo ratios showing *situs inversus* in the F1 generation compared with WT embryos (data not shown). Given that KV is formed from dorsal forerunner cells (DFCs), HSF1 MO was injected into DFCs at the 50% epiboly stage to specifically inhibit the expression of HSF1 in KV. As shown in Figure 2a, DFC injection of MO also resulted in a significant *situs inversus*.

**Figure 2.**
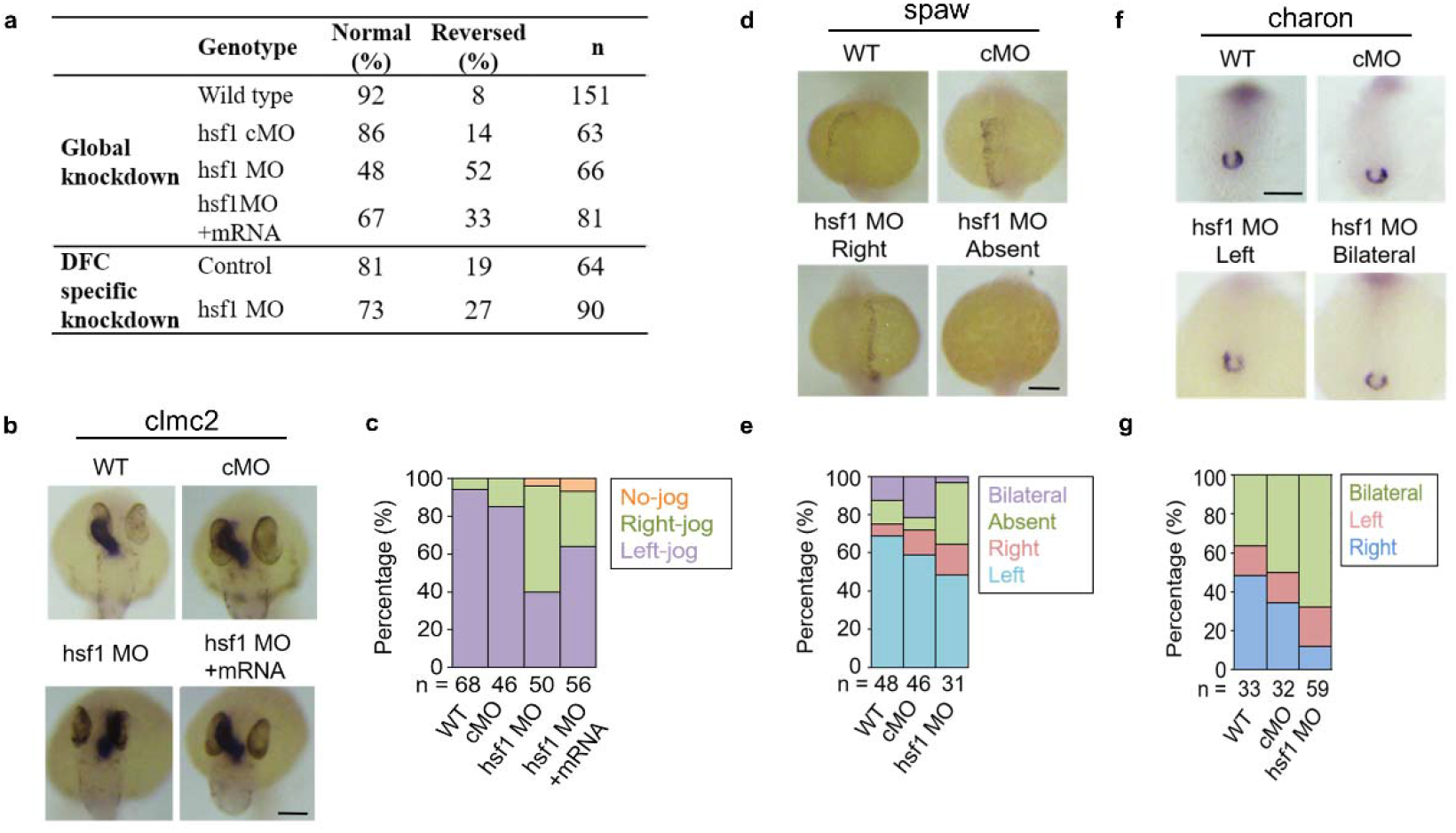
HSF1 is required for the LR asymmetry establishment of zebrafish embryos. (a) The effect of hsf1 knockdown by morpholino (MO) injection on the LR asymmetry establishment in zebrafish embryos. (b) Whole mount in situ hybridization of clmc2 mRNA at 30 hpf in wild type zebrafish embryos (WT) or embryos treated with hsf1 MO or control MO (cMO) with the addition of hsf1 mRNA. Scale bar: 100 μm. (c) Statistical analysis of the results in (b). (d) Whole mount in situ hybridization of spaw mRNA in wild type zebrafish embryos (WT) or embryos treated with hsf1 MO or control morpholino (cMO) at 23-somite stage. (e) Statistical analysis of the results in (d). (f) Whole mount in situ hybridization of charon mRNA in wild type zebrafish embryos (WT) or embryos treated with hsf1 MO or control morpholino (cMO) at 10-somite stage. Scale bars: 100μm. (g) Statistical analysis of the results in (f).

It is believed that the LR axis determination of zebrafish embryos is mediated by Nodal signals^24^. We found that knockdown of HSF1 caused severe disruption of the asymmetric expression of the nodal signaling proteins Spaw and Charon, indicating that asymmetrically activated HSF1 is an early event and could regulate nodal signals during LR axis establishment (Figure 2d-g). Taken together, these results suggest that HSF1 is asymmetrically activated in KV under nodal flow and plays crucial roles in LR symmetry breaking during zebrafish embryonic development.

### 3. Fluid shear stress immediately activates HSF1

As aforementioned, the effect of nodal flow could be mediated by chemical and/or mechanical sensory pathways. Recently, the heat shock response was reported to be induced by compressive loading in bovine caudal disc organ culture^25^. In addition, synovial cells stimulated by fluid shear initiate a heat shock response^26^. In rat cardiac vascular endothelial cells, stretch-sensitive ion channels regulate the activity of HSF1^27^. These studies indicate the possible function of the heat shock response in mechanical stress sensation. Thus, we investigated whether HSF1 activation in the presence of nodal flow is due to mechanical sensation of fluid shear stress. Madin-Darby canine kidney (MDCK) cells, an epithelial cell line originating from the distal renal tubular segment, were subjected to flow shear stress using a flow chamber device for cell culture to mimic fluid shear stress^28,29^ (Figure 3a). Western blot experiments demonstrated that fluid shear stress induced a significant increase in HSF1 phosphorylation on the positive regulatory serine 326 reside and a decrease in the negative regulatory serine 303^30^ residue within 5 minutes (Figure 3b and c). In addition, a heat shock element (HSE)-driving luciferase reporter gene (HSE-Luc) assay also demonstrated elevated transcriptional regulatory activity of HSF1 after fluid shear stress stimuli (Figure 3d).

**Figure 3.**
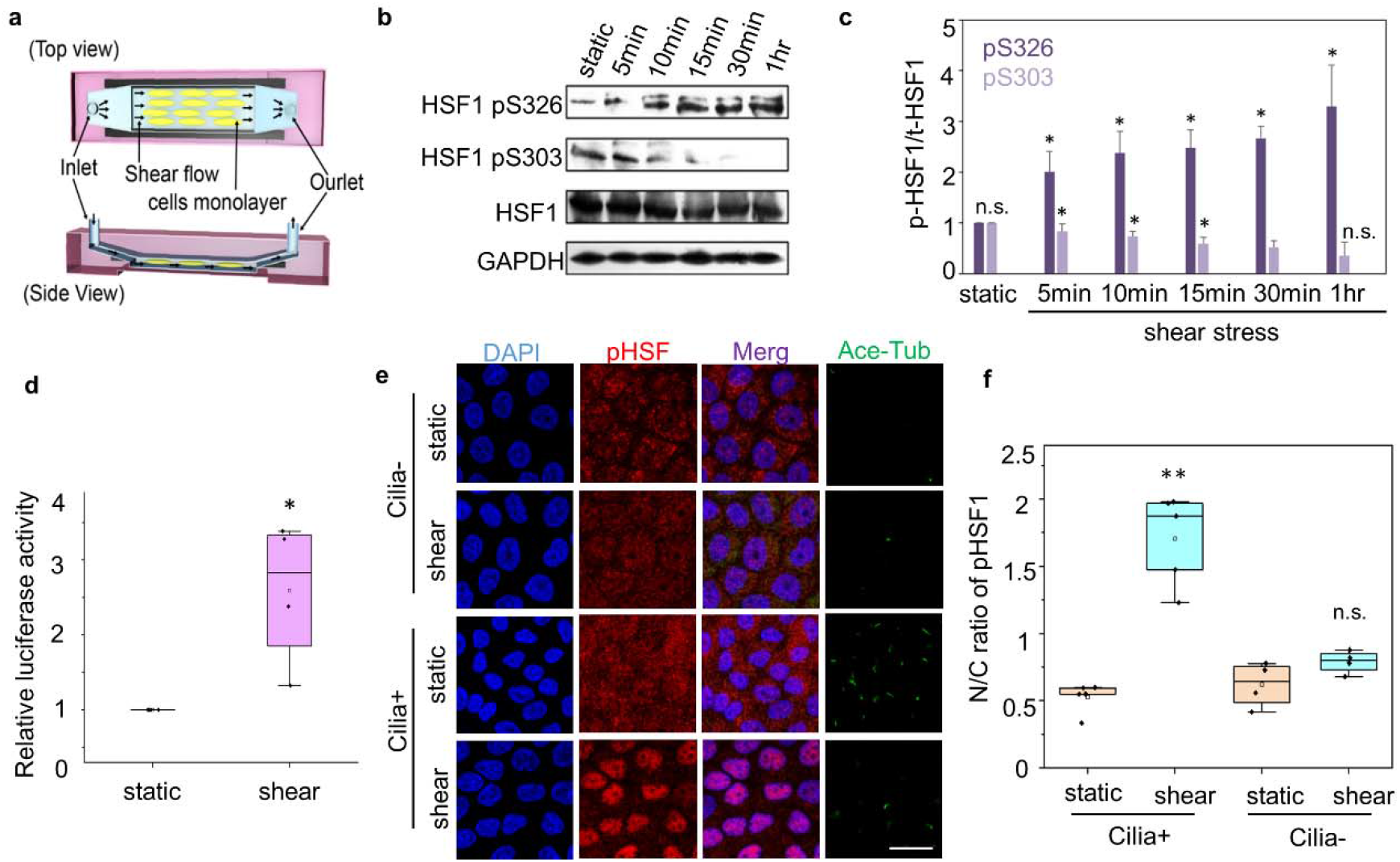
Immediate activation of HSF1 by fluid shear stress. (a) The illustration of the flow chamber device for fluid shear stress experiments. (b) Western blot results of phosphorylated HSF1 at Serine 326 or 303 in MDCK cells after shear stress stimuli for different time. (c) Statistical analysis of the results in (b) (n = 3).The ratio of phosphorylated HSF1 at Serine 326 in nuclei and cytoplasm (N/C) in (b) (n = 10). (d) Luciferase reporter assay for HSE activity in MDCK cells treated with shear stress for 1 h. (n = 3). (e) Fluorescent immunostaining of phosphorylated HSF1 at Serine 326 in MDCK cells 30 min after shear stress. Scale bar: 25μm. (f) The ratio of phosphorylated HSF1 at Serine 326 in nuclei and cytoplasm (N/C) in (e).

To assess the function of cilia in the activation of HSF1 by fluid shear stress, we compared the nuclear transport of HSF1 by immunofluorescence staining in MDCK in the presence or absence of cilia. HSF1 nuclear transport was more obvious in cells with cilia than in those without cilia, indicating the contribution of cilia to the activation of HSF1 under fluid shear stress (Figure 3e and f). These observations are consistent with the mechanical sensing function of cilia, which has been reported in different cell types^31^.

### 4. Mechanical sensitivity of HSF1 is conserved in various types of mechanical stimuli

Next, to evaluate the versatility of HSF1 in mechanical sensing, other types of mechanical stimuli were applied to MDCK cells, including mechanical stretch, hypergravity and substrate stiffness. As shown in Supplemental Figure S1 and S2, mechanical stretch and hypergravity stimuli also triggered rapid nuclear transportation of HSF1 and a significant increase in hsp70 gene expression. Moreover, HSF1 activity was also regulated by substrate stiffness. Compared to the compliant substrate (0.5 kPa), adhesion to the stiff substrate (500 kPa) significantly induced immediate nuclear transportation of HSF1 (Figure 4). To evaluate the conservation of the mechanical sensing function of HSF1 in different cell types, we examined the activation of HSF1 by substrate stiffness in mesenchymal stem cells (MSCs), which are generally used as a cell model for mechanical sensing research^32,33^. The results demonstrated rapid nuclear transportation of HSF1 in MSCs on stiff substrates, which was not observed on soft substrates (Supplemental Figure S3). Collectively, these results suggest that HSF1 is a versatile mechanosensor that can immediately transduce extracellular mechanical stress to gene transcriptional regulation in cells.

**Figure 4.**
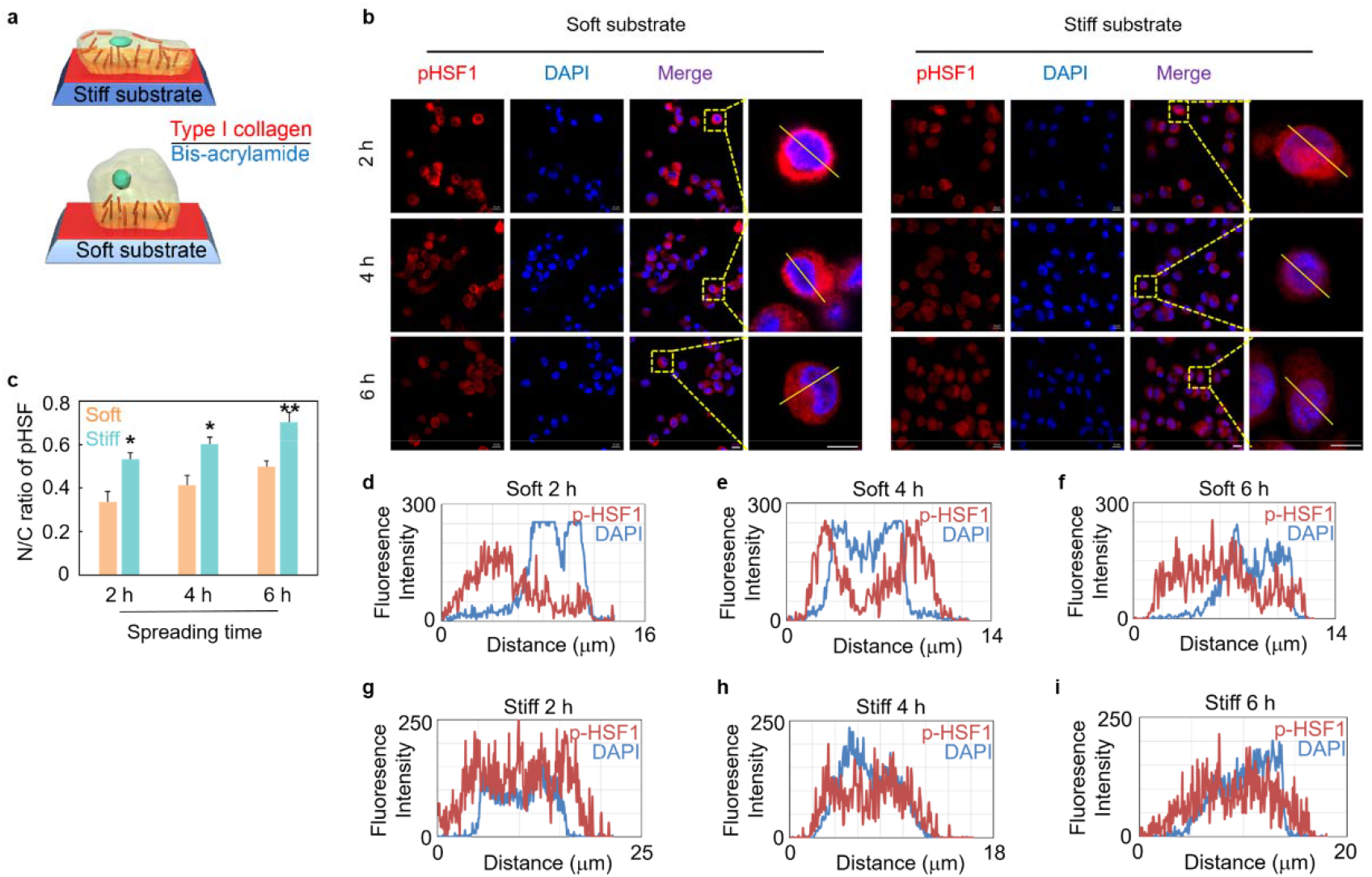
HSF1 activity was regulated by substrate stiffness. (a) The illustration of the experiment device for substrate stiffness stimuli to cultured cells. (b) Fluorescent immunostaining of phosphorylated HSF1 at Serine 326 in MDCK cells under substrate stiffness stimuli. Scale bar: 10μm. (c) The ratio of phosphorylated HSF1 at Serine 326 in nuclei and cytoplasm (N/C) in (b) (n = 9). (d-i) Fluorescence intensity of phosphorylated HSF1 at Serine 326 and DAPI along the lines in the magnified figures in (b).

### 5. Mechanical shear stress activates HSF1 mainly through the Ca^**2+**^**-Akt signaling axis**

To investigate the molecular mechanism underlying HSF1 activation triggered by mechanical stress, we screened signaling pathways that are generally evidenced as important mediators in extracellular signal transduction, including MEK, MAPK, Akt, PKC, and CaMKII^34-38^. The results showed that HSF1 activation under fluid shear stress was strikingly blocked by the Akt inhibitor, whereas a moderate decrease in HSF1 activity was observed in the presence of MAPK, MEK, CaMKII, or PKC inhibitors (Figure 5a and b). These results were consistent with a previous report that Akt could directly interact with HSF1 and phosphorylate it at the positive regulatory residue serine 326^39^.

**Figure 5.**
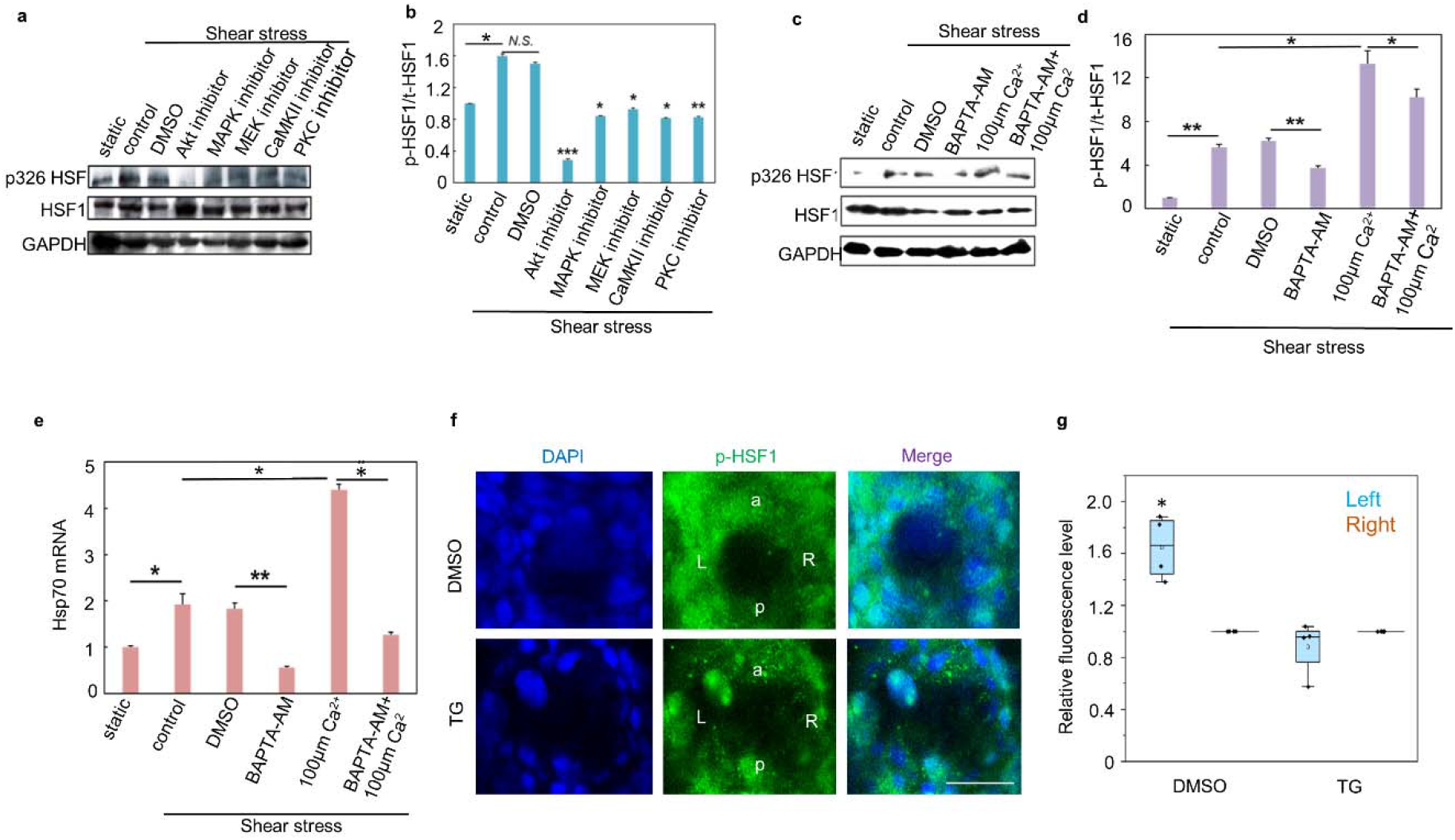
Ca^2+^ and Akt signals were involved in HSF1 activation by shear stress. (a) Western blot results of phosphorylated HSF1 at Serine 326 and total HSF1 in MDCK cells under shear stress in the presence of inhibitors or DMSO. (b) Statistical analysis of the results in (a) (n = 3). (c) Western blot results of phosphorylated HSF1 at Serine 326 and total HSF1 in MDCK cells under shear stress in the presence of Ca2+ antagonist or Ca2+. (d) Statistical analysis of the results in (b) (n = 3). (e) The expression level of hsp70 mRNA in MDCK cells under shear stress in the presence of Ca2+ antagonist or Ca2+ (n = 3). (f) Immunostaining of zebrafish embryos treat with DMSO(DMSO) or embryos treated with 1μM thapsigargin(TG). scale bar:25μm. (g) The ratio of phosphorylated HSF1 at Serine 326 in the left and right of KV in (f) (n = 3).

Many studies have demonstrated that fluid shear stress stimulation induces rapid and temporal intracellular Ca^2+^ increases in many cell types, including MDCK^40-43^. Moreover, Ca^2+^ has a crucial effect on Akt signaling in various cell types^44-47^. Thus, Ca^2+^ signaling may be an upstream signal involved in the immediate activation of HSF1 by mechanical shear stress. To assess this hypothesis, intracellular Ca^2+^ was inhibited by the membrane-permeable Ca^2+^ chelator BAPTA-AM or promoted by additional treatment with Ca^2+^ in the presence of fluid shear stress stimuli. As shown in Figure 5c-e, BAPTA-AM dramatically blocked fluid shear stress-induced HSF1 activation, which could be rescued by excess Ca^2+^. Moreover, additional Ca^2+^ synergistically upregulated HSF1 activity together with fluid shear stress. These results indicate that rapid activation of HSF1 by mechanical stress is mainly mediated by Ca^2+^-Akt signaling pathway. During zebrafish embryonic development, it has been reported that the left-biased Ca^2+^ concentration in KV plays a vital role in LR symmetry breaking^9,10^. In the aforementioned study, we found that Ca^2+^ signaling was crucial for the rapid activation of HSF1 induced by fluid shear stress. To assess whether the left-biased activation of HSF1 in KV also requires Ca^2+^ signaling, we treated zebrafish embryos with an inhibitor of the endoplasmic reticulum Ca-ATPase (thapsigargin, TG) to deplete intracellular Ca^2+^ stores. As shown in Figure 5f and g, the asymmetrical activation of HSF1 was largely blocked by inhibition of Ca^2+^. Thus, we propose a model in which the left-biased activation of HSF1 in KV transduces the mechanical shear stress of nodal flow into gene transcriptional regulation in the dependence of Ca^2+^ signal during LR axis establishment of zebrafish embryos (Supplemental Figure S4a).

## Discussion

Quite a few hypotheses regarding how KV perceives symmetry and regulates symmetry destruction have been proposed in the previous studies, including that cilia perceive the mechanical signal of fluid flow 31,48. Many studies have shown that the distribution of cilia is not uniform and that the flow velocity of the fluid in the cavity caused by cilia is not the same^7^. This makes the mechanical environment in KV different, which provides the possibility for the destruction of KV symmetry caused by cilia sensing the mechanical stimulus. Assuming that the cilia on the left and right sides of the KV produce asymmetric signals, the signal cascade will be initiated, which will lead to asymmetric gene expression. However, this mechanical sensing hypothesis lacks a pivotal mechanism for how the difference in the mechanical microenvironment in KV is transduced into gene expression regulation. In our research, we demonstrated that HSF1 is sensitive to fluid shear stress and immediately transduces mechanical stimuli into gene transcriptional regulation. Thus, our findings support the important role of mechanical conduction in the mechanism of LR symmetry breaking during zebrafish development. According to our results, HSF1 activation is dependent on the cilia, which further confirms the function of cilia in sensing nodal flow.

During zebrafish embryonic development, charon is transiently right-biased expressed in the regions embracing KV at the 8-10 somite stage, which is required for LR patterning^49-52^. Miss-expression of charon elicited phenotypes similar to those of mutant embryos defective in Nodal signaling, which resulted in a loss of LR polarity. We found that HSF1 was asymmetrically activated in KV as early as 8 somite states, and injection of HSF1 MO led to a randomized expression of charon and spaw. Thus, asymmetrical activation of HSF1 may be a relatively early event during LR axis determination. Moreover, using bioinformatics analysis, we found several heat shock elements (HSEs) in the promoter region of zabrafish charon gene (Supplemental Figure S4b). Thus, we hypothesize that HSF1 may contribute to the regulation of charon expression during LR axis establishment. Further studies about this hypothesis are being conducted.

In the response to mechanical stimuli by cells, mechanical signals must be transduced into the nuclei to influence gene expression. Thus, the responses of cells to external mechanical stimuli are time-dependent. However, the time effect of mechanical transduction processes has not been completely studied. In this paper, we identified that HSF1, as a transcription factor, was immediately activated by mechanical stimuli and it rapidly transmitted mechanical signals into gene transcriptional regulation upon various types of mechanical stimuli. Similar to the immediate early genes, the first round of gene transcription after stimuli should be the “gateway” of the complete mechanical response. Thus, the immediate early activation of HSF1 by mechanical stress may be a conserved signal transduction mechanism for gene transcription regulation in the rapid response to mechanical stimuli by cells.

The heat shock response is a fundamental defense against stress. As a highly conserved transcription factor in organisms, HSF1 regulates (induces or represses) the expression of genes that are involved in a wide range of cellular behaviors. Thus, the rapid activation of HSF1 serves as a model for immediate transcriptional responses and has fostered many ground-breaking insights into the regulatory mechanisms of gene expression. Our findings unveiled that in addition to thermal and oxidative stresses, HSF1 also responds to mechanical stresses, which is important for gene regulation triggered by mechanical clues during embryonic development and other physiological and pathological transformations.

## Materials and Methods

### Cell and fish maintenance

MDCK (Madin-Darby canine kidney) cells and human mesenchymal stem cells were grown in DMEM culture medium supplemented with 10% fetal bovine serum (FBS) (Gibco), 100 mg/ml streptomycin and 100 units/ml penicillin (Gibco) in a humidified incubator at 37 °C and with 5% CO_2_ atmosphere. All zebrafish husbandry was performed under standard conditions in accordance with institutional and national ethical and animal welfare guidelines. Collected embryos were grown at 28.5 °C.

### Antibodies and reagents

Antibodies against HSF1, anti-HSF1 (phospho S326) and anti-HSF1 (phospho S303) were purchased from Abcam. Monoclonal anti-Acetylated Tubulin antibody was purchased from Sigma. Alexa Fluor 546-labeled goat anti-rabbit secondary polyclonal antibody and Alexa Fluor 488-labeled goat anti-mouse secondary polyclonal antibody were purchased from Life Technology. Monoclonal antibody against GAPDH, horseradish peroxidase (HRP)-coupled goat anti-rabbit secondary polyclonal antibody and horseradish peroxidase (HRP)-coupled goat anti-mouse secondary polyclonal antibody were obtained from ZSGB-BIO.

Inhibitors of Akt (MK-2206 2HCl), MAPK (SB203580), MEK (U0126), CaMKII (KN-93 Phosphate), PKC (Staurosporine) and Ca^2+^ (BAPTA-AM), were purchased from Selleck.

### CRISPR/Cas9-Mediated Knockdown in Zebrafifish

CRISPR/Cas9-mediated knockdown zebrafifish were obtained from the China Zebrafish Resource Center (CZRC). Wild-type AB/TU and mutant zebrafish were raised and maintained under standard conditions. Zebrafish embryos were obtained by artificial insemination and reared at 28 °C. All animal experiments were conducted in accordance with the Guiding Principles for the Care and Use of Laboratory Animals and were approved by the Institute of Hydrobiology, Chinese Academy of Sciences.

To study the function of gene HSF1, we employed the Cas9/gRNA system to generate NOD1 knockout zebrafish. A gRNA with 20-bp “target sequence” was designed, which starts with two GG residues at the 5′ end for efficient transcription from the T7 promoter and ends with the protospacer adjacent motif (PAM) NGG at the 3′ end, which is indispensable for Cas9 binding and cleavage. Cas9 mRNA and gene HSF1 gRNA were microinjected into one-cell embryos of zebrafish. The results from sequencing of PCR fragments from a single zebrafish about 2 months old revealed two or more peaks at the same location. As expected, the Cas9/gRNA-mediated mutations occurred at or near the target site.Two genotypes of zebrafish CZ1196/ihb287; CZ1203/ihb294: one is between 412bp to 421bp of the wild-type hsf1 coding sequence, TGGCCAGTTT, is deleted. The mutated hsf1 codes for a truncated protein containing 152 aa, 133 aa of which is identical to wildtype hsf1. Anothor is between 418 bp to 422 bp of the wild-type hsf1 coding sequence, GTTTT, is deleted. The mutated hsf1 codes for a truncated protein containing 233 aa, 139 aa of which is identical to wildtype hsf1.

### Mechanical stimuli apparatuses

#### Flow chamber

A rectangular parallel plate perfusion chamber, designed and presented by professor Jeng-Jiann Chiu Lab in the National Health Research Institutes of Taiwan, was used for fluid shear stress exposure. This system comprises a transparent polymethylmethacrylate plate, two silastic rubber gaskets, and a standard glass coverslip. The coverslip with near 90% confluent monolayer of MDCK cells was mounted over the groove with the cells facing the inside, and an approximate 200 μm high gap was formed over the MDCK cells. Then we collected the treated samples from each condition for the following analyses.

#### Mechanical stretch

PDMS curing agent (sylgard184; Dow corning Corp) was mixed with base agent in a mass ratio of 10:1 in 50 ml centrifugal tubes, and centrifuged in a bench-top centrifuge to remove air bubbles, which was subsequently cast onto a single-well Petri dish (35 × 10 mm, Corning) and cured at 65 °C for 12 h. The PDMS substrates were oxidized in an oxygen plasma cleaner, which generated silanol group (Si-OH) on the surface of the PDMS, then sterilized by UV radiation for 2 h, coated with fibronectin and left in a laminar flow cabinet for at least 2 h. Then, MDCK cells achieving 90% confluence were subjected to mechanical stress with the Cyclic Stress Unit. The unit was placed in a humidified incubator with 5% CO_2_ at 37 °C. Cyclic deformation (20 cycles/min) and up to 10% elongation was applied. Then we collected the treated samples from each condition for the following analyses.

#### Hypergravity stress

MDCK Cells (5 × 10^4^) were placed in an incubator at 37 °C with 40 ×g with hypergravity treatments using a centrifuge. Samples were collected at 30 min, 1 h, 2 h and 4 h. Then, the treated samples from each condition were used for the following analyses.

#### Substrate stiffness

Polyacrylamide gels with variable Young’s moduli were prepared as described previously^33^. Briefly, acrylamide and bis-acrylamide mixture with indicated concentrations was allowed to polymerize on a glass slide. Then, the gel was covered by sulfosuccinimidyl-6-[4′-azido-2′-nitrophenylamino] hexanoate (Sulfo-SANPAH; Pierce).

After exposure to UV light for 10 min twice, the polyacrylamide sheet was washed twice and incubated with a solution of type I collagen (0.2 mg/ml; Gibco BRL) overnight at 4°C.

#### Immunofluorescence staining and confocal microscopy

MDCK cells were fixed with 4% paraformaldehyde (PFA), permeabilized with 0.5% Triton X-100, blocked with 5% BSA for 1 h at room temperature. Then, cells were probed with antibodies overnight, followed by incubation with Alexa Fluor 546-labeled goat anti-rabbit secondary polyclonal antibody and Alexa Fluor 488-labeled goat anti-mouse secondary polyclonal antibody for 1 h. After washing with PBST for several times, cell nuclei were stained by DAPI. Then, the fluorescent images were captured with a ×63 objective mounted on a LSM710 confocal microscope.

The zebrafish embryos were fixed in BT fixative buffer [4% paraformaldehyde, 0.15 mM CaCl_2_, 4% sucrose in 0.1 M PBS (pH 7.3)] overnight at 4 °C. After being rinsed 3-5 times with PBST, embryos were blocked at room temperature for 1 h in 5% BSA and 1% DMSO in PBST. Embryos were then incubated overnight at 4 °C with antibodies. After washing with PBST three times, embryos were incubated overnight at 4°C with Alexa Fluor 488-conjugated AffiniPure Donkey anti-mouse IgG and Alexa Fluor 546-conjugated AffiniPure Goat anti-rabbit IgG. Embryos were then washed with PBST and stained by DAPI. Then, embryos were transferred into an anti-fade reagent and stored at 4 °C for up to 3 d. Immunostained embryos were imaged under a Zeiss 710META laser

scanning confocal microscope with a 63×objective. A KV/cilia image was a sum of multiple focal planes (z-series), combined using ImageJ. The fluorescent intensity was analyzed using Image-Pro Plus 6.0.

#### Quantitative Real-time PCR (qRT-PCR)

The qRT-PCR was conducted as described previously^53^. Total RNA was isolated using the Trizol reagent (Invitrogen). For qRT-PCR analysis, it was performed in accordance with the manufacturer’s instructions using FastQuant RT Kit (With gDNase) (Tiangen). Relative mRNA levels of each gene were analyzed according to the comparative Ct method and were normalized to the expression of GAPDH. The following sets of primers were used in the PCR amplification:

hsp70 (forward, 5□-AGCTGGAGCAGGTGTGTAAC-3□; reverse, 5□-GGGGAAGAA GTCCTAATCCACC-3□). GAPDH (forward, 5□-AGTCAACGGATTTGGCCGTA-3□; reverse, 5□-CCGTTCTCAGCCTTGACTGT-3□).

For all the qRT-PCR experiments, the correlation coefficient and amplification efficiency in the reaction using each primer are 0.97–1.02 and more than 97.9%, respectively.

#### MO oligonucleotides injections

For totally knockdown, morpholinos (Gene Tools) were injected into the yolk of one-cell stage embryos. DFCs specific knockdown was performed as described previously^54^. Briefly, the morpholino was mixed with rhodamine (Sigma) and injected into the yolk at the 512-cell stage. Then, at 60-70% epiboly stages, the embryos with red fluorescence in DFCs were sorted and raised for further investigations.

#### Double-luciferase report gene experiment

RL(500ng/ul,clontch)and pHSE-luc(500ng/ul,clontch) were cotransfected into MDCK cell line using FuGENE HD for 24h. we use shear stress stimulate cell 1h after transfection. The cells were lysed using 1X passive lysate. Firefly luciferase was used as a transfection control group, and the luciferase activity was measured using a dual luciferase reporter assay system.

#### Western blotting

Western blotting was conducted as described previously^33^. Proteins (40 μg) from cell homogenates were separated on a 10% SDS-polyacrylamide gel and electrotransfered to polyvinylidene fluoride membranes (Millipore). Each membrane was washed with Tris-buffered saline Tween-20, blocked with 5% skim milk power for 1 h, and incubated with appropriate primary antibodies at dilutions recommended by the supplier. Membranes were washed in TBST and probed with an appropriate HRP conjugated secondary antibody. The oxidase components were detected by chemiluminescence (Super ECL Plus, Huaxingbio Science).

#### Whole-mount in situ hybridization

Whole-mount in situ hybridization analyses were performed as described previously^24^. The primers used for probes synthesis are as follows: southpaw, CCGCTGTACATGATGCAGTT and GTAAGCGTGGTTTGTTGGGT; charon, AAGACTTTGAATCCTCCGGG and ACGTTTCTGTTTGCAGGGAC; cmlc2, TTGTGCAGTTATCAGGGCTC and TTAACAGTCTGTAGGGGGCA; hsp70, GAAGCTTCTGGAGGATTTCT and CAATGTGGAGGACATGCACG.

## H2: Supplementary Materials

Fig. S1. Immediate activation of HSF1 by mechanical stretch.

Fig. S2. Immediate activation of HSF1 by hypergravity stimuli.

Fig. S3. HSF1 activity in MSCs was regulated by substrate stiffness.

Fig. S4. Pattern diagram and charon promoter analysis.

## General

We thank A.M. Meng (Tsinghua University) for technical assistance and discussions in the course of the preparation of this manuscript. We also thank H. Yamaguchi (The University of Tokyo) for technical assistance of zebrafish experiments. We wish to thank Dr. XunWei Xie from the China Zebrafish Resource Center for helping in the generation of geneX-mutant zebrafish.

## Author contributions

Y.B.F., H.T., D.Y.C. and J.D. designed the study and interpreted experiments. J.D., S.K.L., and L.Y.G performed cell and zebrafish experiments. Z.G., T.K., A.S., and Q.P.Z. helped with the zebrafish experiments. J.F.Y., L.G., L.S.Z., Y.Y.L., and X.Q.F. helped with the cell mechanical stimuli experiments. Y.B.F., H.T., D.Y.C. and J.D. conceived and supervised this project and wrote the paper.

## Supplementary Materials

**Figure S1.**
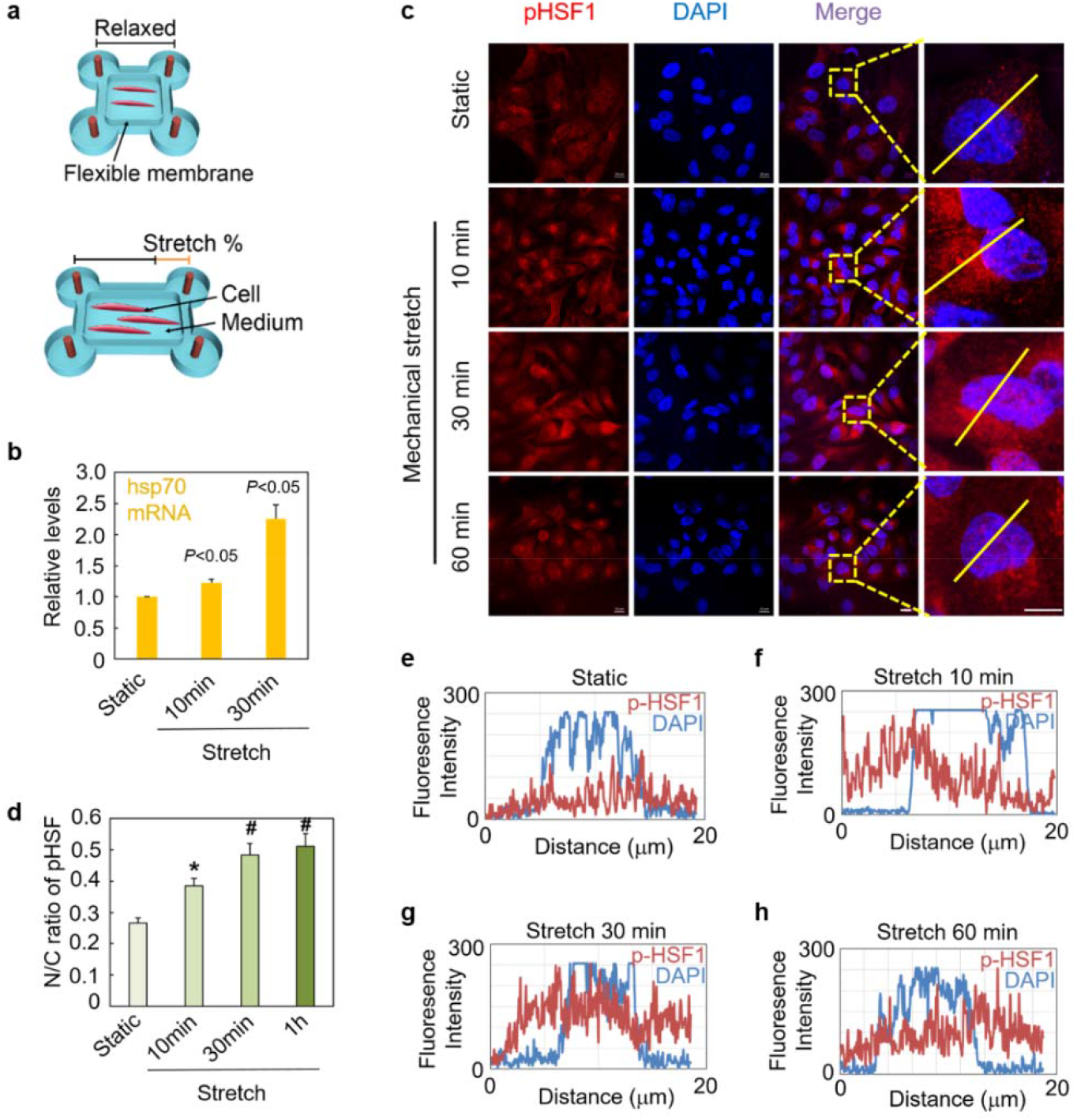
Immediate activation of HSF1 by mechanical stretch. (a) The illustration of the experiment device for mechanical stretch applied to cultured cells. (b) The expression level of hsp70 mRNA in MDCK cells after mechanical stretch stimuli for 10 min or 30 min measured by qRT-PCR (n = 3). (c) Fluorescent immunostaining of phosphorylated HSF1 at Serine 326 in MDCK cells under mechanical stretch. Scale bar: 10 μm. (d) The ratio of phosphorylated HSF1 at Serine 326 in nuclei and cytoplasm (N/C) in (c) (n = 16). (e-h) Fluorescence intensity of phosphorylated HSF1 at Serine 326 and DAPI along the lines in the magnified figures in (c).

**Figure S2.**
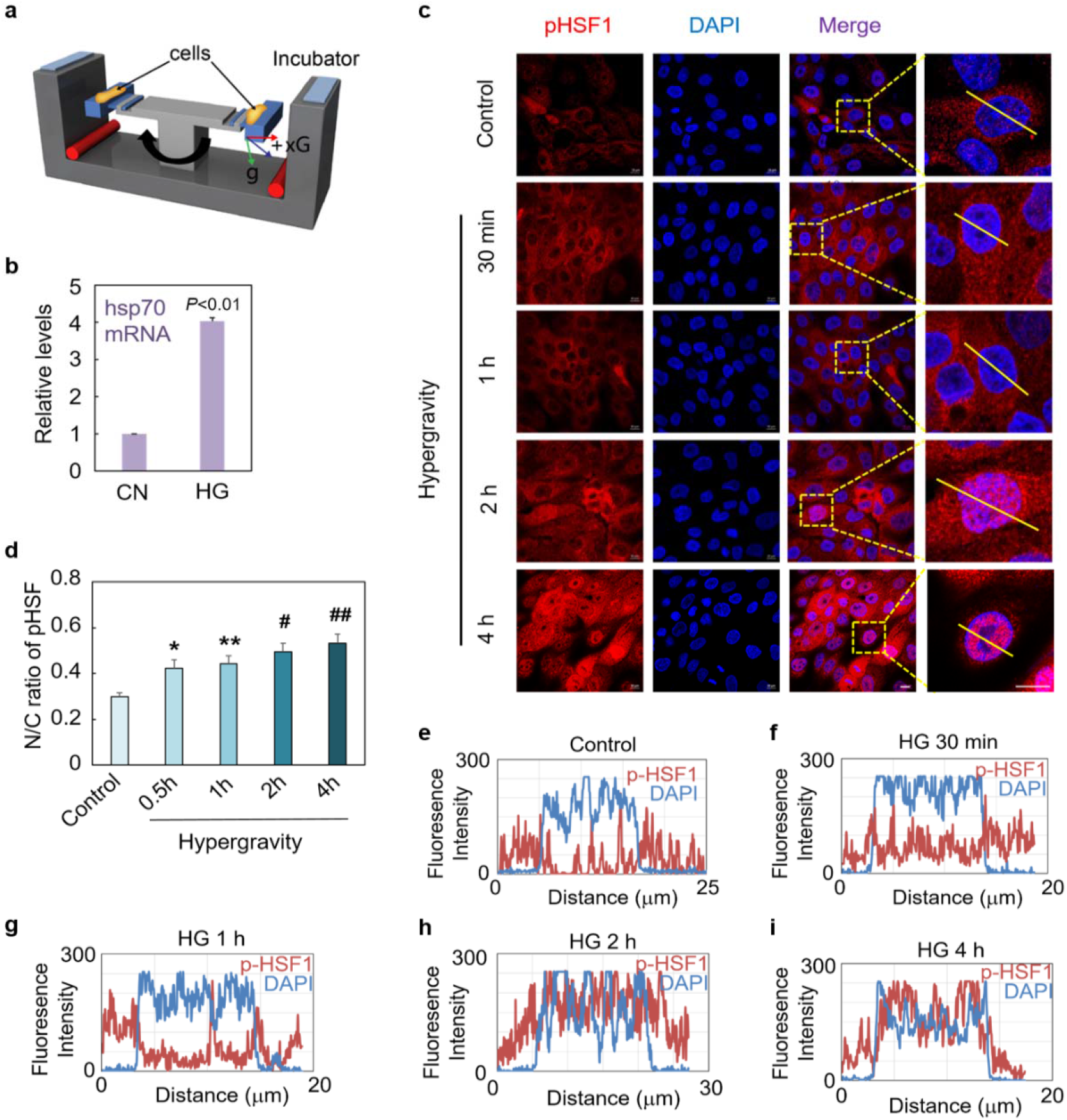
Immediate activation of HSF1 by hypergravity stimuli. (a) The illustration of the experiment device for mechanical stretch applied to cultured cells. (b) The expression level of hsp70 mRNA in MDCK cells after mechanical stretch stimuli for 10 min or 30 min measured by qRT-PCR (n = 3). (c) Fluorescent immunostaining of phosphorylated HSF1 at Serine 326 in MDCK cells under mechanical stretch. Scale bar: 10 μm. (d) The ratio of phosphorylated HSF1 at Serine 326 in nuclei and cytoplasm (N/C) in (c) (n = 16). (e-i) Fluorescence intensity of phosphorylated HSF1 at Serine 326 and DAPI along the lines in the magnified figures in (c).

**Figure S3.**
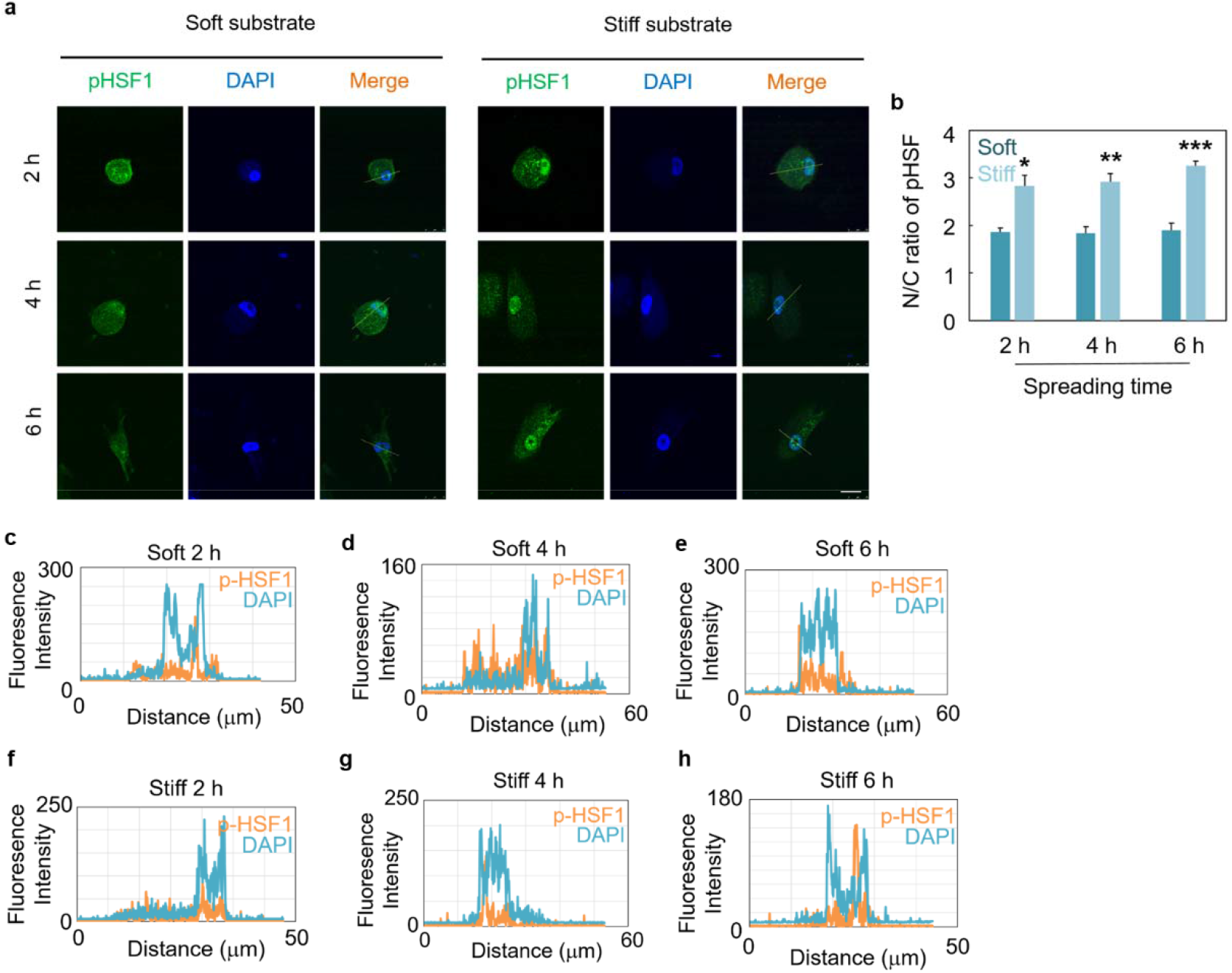
HSF1 activity in MSCs was regulated by substrate stiffness. (a) Fluorescent immunostaining of phosphorylated HSF1 at Serine 326 in MSCs under substrate stiffness stimuli. Scale bar: 25 μm. (b) The ratio of phosphorylated HSF1 at Serine 326 in nuclei and cytoplasm (N/C) in (a) (n = 9). (c-h) Fluorescence intensity of phosphorylated HSF1 at Serine 326 and DAPI along the lines in the magnified figures in (b).

**Figure S4.**
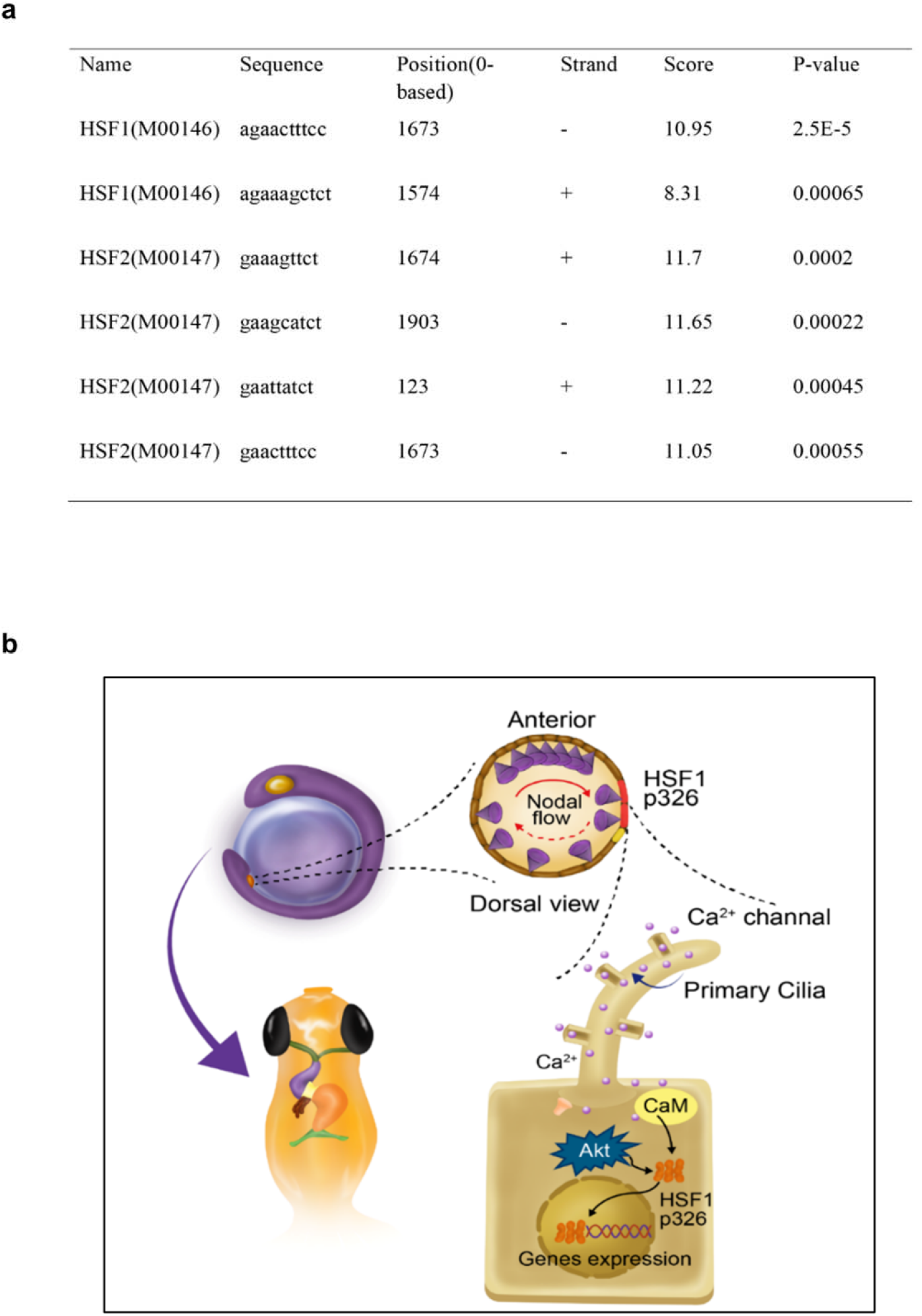
Pattern diagram and charon promoter analysis. (a) Illustration of gene transcriptional regulation by HSF1 under nodal flow shear stress during LR asymmetry establishment of zebrafish embryos. (b) Zebrafish charon promoter (−2 kb) binding site analysis was performed using integrated Web tools.

## References

1 Freund, J. B., Goetz, J. G., Hill, K. L. & Vermot, J. Fluid flows and forces in development: functions, features and biophysical principles. Development 139, 1229–1245 (2012).

2 Hojo, M. et al. RightlJelevated expression of charon is regulated by fluid flow in medaka Kupffer’s vesicle. Development, growth & differentiation 49, 395–405 (2007).

3 Kreiling, J. A., Williams, G. & Creton, R. Analysis of Kupffer’s vesicle in zebrafish embryos using a cave automated virtual environment. Developmental Dynamics 236, 1963–1969 (2007).

4 Cartwright, J. H., Piro, O. & Tuval, I. Fluid-dynamical basis of the embryonic development of left-right asymmetry in vertebrates. Proc Natl Acad Sci U S A 101, 7234–7239, doi:10.1073/pnas.0402001101 (2004).

5 Shah, A. S., Ben-Shahar, Y., Moninger, T. O., Kline, J. N. & Welsh, M. J. Motile cilia of human airway epithelia are chemosensory. Science 325, 1131–1134, doi:10.1126/science.1173869 (2009).

6 Yoder, B. K. Role of primary cilia in the pathogenesis of polycystic kidney disease. J Am Soc Nephrol 18, 1381–1388, doi:10.1681/ASN.2006111215 (2007).

7 Montenegro-Johnson, T. D., Baker, D. I., Smith, D. J. & Lopes, S. S. Three-dimensional flow in Kupffer’s Vesicle. J Math Biol 73, 705–725, doi:10.1007/s00285-016-0967-7 (2016).

8 Sampaio, P. et al. Left-right organizer flow dynamics: how much cilia activity reliably yields laterality? Dev Cell 29, 716–728, doi:10.1016/j.devcel.2014.04.030 (2014).

9 Yuan, S., Zhao, L., Brueckner, M. & Sun, Z. Intraciliary calcium oscillations initiate vertebrate left-right asymmetry. Current Biology 25, 556–567 (2015).

10 Takao, D. et al. Asymmetric distribution of dynamic calcium signals in the node of mouse embryo during left–right axis formation. Developmental biology 376, 23–30 (2013).

11 Voellmy, R. & Boellmann, F. in Molecular Aspects of the Stress Response: Chaperones, Membranes and Networks 89–99 (Springer, 2007).

12 Vígh, L. et al. in Molecular Aspects of the Stress Response: Chaperones, Membranes and Networks 114–131 (Springer, 2007).

13 Ryno, L. M. et al. Characterizing the altered cellular proteome induced by the stress-independent activation of heat shock factor 1. ACS chemical biology 9, 1273–1283 (2014).

14 Kus-Liskiewicz, M. et al. Impact of heat shock transcription factor 1 on global gene expression profiles in cells which induce either cytoprotective or pro-apoptotic response following hyperthermia. BMC genomics 14, 456, doi:10.1186/1471-2164-14-456 (2013).

15 Hahn, J. S., Hu, Z., Thiele, D. J. & Iyer, V. R. Genome-wide analysis of the biology of stress responses through heat shock transcription factor. Molecular and cellular biology 24, 5249–5256, doi:10.1128/MCB.24.12.5249-5256.2004 (2004).

16 Page, T. J. et al. Genome-wide analysis of human HSF1 signaling reveals a transcriptional program linked to cellular adaptation and survival. Molecular bioSystems 2, 627–639, doi:10.1039/b606129j (2006).

17 Birch-Machin, I. et al. Genomic analysis of heat-shock factor targets in Drosophila. Genome biology 6, R63, doi:10.1186/gb-2005-6-7-r63 (2005).

18 Ishikawa, T., Igarashi, T., Hata, K. & Fujita, T. c-fos induction by heat, arsenite, and cadmium is mediated by a heat shock element in its promoter. Biochem Biophys Res Commun 254, 566–571, doi:10.1006/bbrc.1998.9979 (1999).

19 Sawai, M., Ishikawa, Y., Ota, A. & Sakurai, H. The proto-oncogene JUN is a target of the heat shock transcription factor HSF1. FEBS J 280, 6672–6680, doi:10.1111/febs.12570 (2013).

20 Wang, G. et al. Suppression of heat shock transcription factor HSF1 in zebrafish causes heat-induced apoptosis. Genesis 30, 195–197 (2001).

21 Evans, T. G., Belak, Z., Ovsenek, N. & Krone, P. H. Heat shock factor 1 is required for constitutive Hsp70 expression and normal lens development in embryonic zebrafish. Comp Biochem Physiol A Mol Integr Physiol 146, 131–140, doi:10.1016/j.cbpa.2006.09.023 (2007).

22 Chou, S. D., Prince, T., Gong, J. & Calderwood, S. K. mTOR is essential for the proteotoxic stress response, HSF1 activation and heat shock protein synthesis. PLoS One 7, e39679, doi:10.1371/journal.pone.0039679 (2012).

23 Jaffe, K. M. et al. c21orf59/kurly Controls Both Cilia Motility and Polarization. Cell Rep 14, 1841–1849, doi:10.1016/j.celrep.2016.01.069 (2016).

24 Koshida, S., Shinya, M., Mizuno, T., Kuroiwa, A. & Takeda, H. Initial anteroposterior pattern of the zebrafish central nervous system is determined by differential competence of the epiblast. Development 125, 1957–1966 (1998).

25 Chooi, W. H., Chan, S. C., Gantenbein, B. & Chan, B. P. Loading-Induced Heat-Shock Response in Bovine Intervertebral Disc Organ Culture. PLoS One 11, e0161615, doi:10.1371/journal.pone.0161615 (2016).

26 Schett, G. et al. Enhanced expression of heat shock protein 70 (hsp70) and heat shock factor 1 (HSF1) activation in rheumatoid arthritis synovial tissue. Differential regulation of hsp70 expression and hsf1 activation in synovial fibroblasts by proinflammatory cytokines, shear stress, and antiinflammatory drugs. J Clin Invest 102, 302–311, doi:10.1172/JCI2465 (1998).

27 Chang, J., Wasser, J. S., Cornelussen, R. N. & Knowlton, A. A. Activation of heat-shock factor by stretch-activated channels in rat hearts. Circulation 104, 209–214 (2001).

28 Duan, Y., Weinstein, A. M., Weinbaum, S. & Wang, T. Shear stress-induced changes of membrane transporter localization and expression in mouse proximal tubule cells. Proc Natl Acad Sci U S A 107, 21860–21865, doi:10.1073/pnas.1015751107 (2010).

29 Tarran, R. et al. Normal and cystic fibrosis airway surface liquid homeostasis. The effects of phasic shear stress and viral infections. The Journal of biological chemistry 280, 35751–35759, doi:10.1074/jbc.M505832200 (2005).

30 Guettouche, T., Boellmann, F., Lane, W. S. & Voellmy, R. Analysis of phosphorylation of human heat shock factor 1 in cells experiencing a stress. BMC Biochem 6, 4, doi:10.1186/1471-2091-6-4 (2005).

31 Ferreira, R. R., Fukui, H., Chow, R., Vilfan, A. & Vermot, J. The cilium as a force sensor-myth versus reality. J Cell Sci 132, doi:ARTN jcs213496 10.1242/jcs.213496 (2019).

32 Engler, A. J., Sen, S., Sweeney, H. L. & Discher, D. E. Matrix elasticity directs stem cell lineage specification. Cell 126, 677–689, doi:10.1016/j.cell.2006.06.044 (2006).

33 Du, J. et al. Integrin activation and internalization on soft ECM as a mechanism of induction of stem cell differentiation by ECM elasticity. Proc Natl Acad Sci U S A 108, 9466–9471, doi:10.1073/pnas.1106467108 (2011).

34 Leevers, S. J. & Marshall, C. J. Activation of extracellular signal-regulated kinase, ERK2, by p21ras oncoprotein. EMBO J 11, 569–574 (1992).

35 Cargnello, M. & Roux, P. P. Activation and function of the MAPKs and their substrates, the MAPK-activated protein kinases. Microbiology and molecular biology reviews : MMBR 75, 50–83, doi:10.1128/MMBR.00031-10 (2011).

36 Dudek, H. et al. Regulation of neuronal survival by the serine-threonine protein kinase Akt. Science 275, 661–665 (1997).

37 Reyland, M. E. Protein kinase C isoforms: Multi-functional regulators of cell life and death. Frontiers in bioscience 14, 2386–2399 (2009).

38 Hudmon, A. & Schulman, H. Neuronal CA2+/calmodulin-dependent protein kinase II: the role of structure and autoregulation in cellular function. Annual review of biochemistry 71, 473–510, doi:10.1146/annurev.biochem.71.110601.135410 (2002).

39 Carpenter, R. L., Paw, I., Dewhirst, M. W. & Lo, H. W. Akt phosphorylates and activates HSF-1 independent of heat shock, leading to Slug overexpression and epithelial-mesenchymal transition (EMT) of HER2-overexpressing breast cancer cells. Oncogene 34, 546–557, doi:10.1038/onc.2013.582 (2015).

40 Praetorius, H. A. & Spring, K. R. Bending the MDCK cell primary cilium increases intracellular calcium. The Journal of membrane biology 184, 71–79 (2001).

41 Yamamoto, K., Korenaga, R., Kamiya, A. & Ando, J. Fluid shear stress activates Ca(2+) influx into human endothelial cells via P2X4 purinoceptors. Circulation research 87, 385–391 (2000).

42 Jafarnejad, M. et al. Measurement of shear stress-mediated intracellular calcium dynamics in human dermal lymphatic endothelial cells. American journal of physiology. Heart and circulatory physiology 308, H697–706, doi:10.1152/ajpheart.00744.2014 (2015).

43 Schwarz, G., Callewaert, G., Droogmans, G. & Nilius, B. Shear stress-induced calcium transients in endothelial cells from human umbilical cord veins. The Journal of physiology 458, 527–538 (1992).

44 Danciu, T. E., Adam, R. M., Naruse, K., Freeman, M. R. & Hauschka, P. V. Calcium regulates the PI3K-Akt pathway in stretched osteoblasts. FEBS letters 536, 193–197 (2003).

45 Nicholson-Fish, J. C., Cousin, M. A. & Smillie, K. J. Phosphatidylinositol 3-Kinase Couples Localised Calcium Influx to Activation of Akt in Central Nerve Terminals. Neurochemical research 41, 534–543, doi:10.1007/s11064-015-1663-5 (2016).

46 Perez-Garcia, M. J. et al. Glial cell line-derived neurotrophic factor increases intracellular calcium concentration. Role of calcium/calmodulin in the activation of the phosphatidylinositol 3-kinase pathway. The Journal of biological chemistry 279, 6132–6142, doi:10.1074/jbc.M308367200 (2004).

47 Divolis, G., Mavroeidi, P., Mavrofrydi, O. & Papazafiri, P. Differential effects of calcium on PI3K-Akt and HIF-1alpha survival pathways. Cell biology and toxicology 32, 437–449, doi:10.1007/s10565-016-9345-x (2016).

48 Omori, T., Winter, K., Shinohara, K., Hamada, H. & Ishikawa, T. Simulation of the nodal flow of mutant embryos with a small number of cilia: comparison of mechanosensing and vesicle transport hypotheses. Roy Soc Open Sci 5, doi:ARTN 18060110.1098/rsos.180601 (2018).

49 Hojo, M. et al. Right-elevated expression of charon is regulated by fluid flow in medaka Kupffer’s vesicle. Development, growth & differentiation 49, 395–405, doi:10.1111/j.1440-169X.2007.00937.x (2007).

50 Schneider, I. et al. Zebrafish Nkd1 promotes Dvl degradation and is required for left-right patterning. Developmental biology 348, 22–33, doi:10.1016/j.ydbio.2010.08.040 (2010).

51 Kawasumi, A. et al. Left-right asymmetry in the level of active Nodal protein produced in the node is translated into left-right asymmetry in the lateral plate of mouse embryos. Dev Biol 353, 321–330, doi:10.1016/j.ydbio.2011.03.009 (2011).

52 Nakamura, T. et al. Fluid flow and interlinked feedback loops establish left-right asymmetric decay of Cerl2 mRNA. Nature communications 3, 1322, doi:10.1038/ncomms2319 (2012).

53 Du, J. et al. Extracellular matrix stiffness dictates Wnt expression through integrin pathway. Sci Rep 6, 20395, doi:10.1038/srep20395 (2016).

54 Amack, J. D. & Yost, H. J. The T box transcription factor no tail in ciliated cells controls zebrafish left-right asymmetry. Curr Biol 14, 685–690, doi:10.1016/j.cub.2004.04.002 (2004).

